# Expanding Threat of Carbapenemase-Producing *Escherichia coli* and *Klebsiella pneumoniae* in Peru: Genomic and Phenotypic Evidence of High-Risk Clones Dissemination

**DOI:** 10.64898/2026.04.28.721358

**Authors:** Edgar Gonzales Escalante, Roky G. Champi-Merino, Luis Alvarado, Juan Carlos Gómez-de-la-Torre, Elizabeth Sierra Chávez, Roxana Sandoval-Ahumada, Giancarlo Pérez Lazo, Casanova Janet, Marco Matta-Chuquisapon, Liliana Violeta Morales Castillo, Escobar Valdiviezo Alejandra, Gonzales-Rodriguez Arturo

## Abstract

Carbapenemase-producing *Enterobacterales* represent a growing global threat due to their extensive antimicrobial resistance and rapid dissemination. This study characterized the phenotypic and genomic features of *Escherichia coli* and *Klebsiella pneumoniae* isolates collected between 2020 and 2022 from four healthcare institutions in Lima, Peru. A total of 320 non-redundant isolates (61 *E. coli* and 259 *K. pneumoniae*) were analyzed through antimicrobial susceptibility testing, polymerase chain reaction, and whole-genome sequencing. The most frequent carbapenemase gene was *bla*_NDM_ (69%), followed by *bla*_KPC_ (16.9%) and *bla_OXA-48-like_* (4.6%). Eleven *K. pneumoniae* isolates co-produced NDM and KPC, and one *E. coli* isolate co-harbored NDM and OXA-48-like. All isolates were multidrug resistant, and 5% were pandrug resistant. Novel β-lactam/β-lactamase inhibitor combinations such as aztreonam/avibactam and cefiderocol showed complete activity against all classes of carbapenemases. Genomic analysis revealed predominant *E. coli* sequence types ST167 and ST410 and *K. pneumoniae* lineages ST147, ST15, ST45, and ST273. The *bla*_NDM-5_ allele was detected for the first time in Peru, mostly in *E. coli ST167*, carried on multireplicon IncF-type plasmids. In *K. pneumoniae*, ST147 was identified as a dominant clone associated with *bla*_NDM-1_, indicating sustained local dissemination of high-risk clonal groups. The coexistence of multiple carbapenemases and plasmid backbones highlights the ongoing evolution of resistance mechanisms. These findings provide actionable evidence to guide treatment strategies in settings with high prevalence of metallo-β-lactamases and underscores the need for continuous genomic surveillance and antimicrobial stewardship to mitigate their clinical and epidemiological impact.

## INTRODUCTION

The increasing prevalence and dissemination of carbapenemase-producing *Enterobacterales* (CPE) represent a major public health concern, as treatment options are severely limited and often involve high-cost antimicrobial agents (1). Consequently, the World Health Organization (WHO) has classified CPE among the highest-priority pathogens, emphasizing the urgent need to monitor their emergence and implement strategies to mitigate this growing global threat (1).

Carbapenemases play a fundamental role in carbapenem resistance, representing the main mechanism of resistance among *Enterobacterales*. These enzymes are classified into Ambler classes A, B, and D, with KPC, NDM, and OXA-48-like variants being the most prevalent worldwide (2). The increasing reports of carbapenemase-producing microorganisms have led to endemic dissemination across Latin America, particularly in countries such as Brazil, Colombia, Argentina, and Mexico, which account for the highest number of publications on this topic (3). In Peru, class A, B, and D carbapenemases have been identified; however, NDM remains by far the most frequently reported (4).

*Enterobacterales* such as *Escherichia coli* and *Klebsiella pneumoniae* are both common human pathogens and asymptomatic colonizers of the gastrointestinal tract and various environmental niches (5,6). *K. pneumoniae* and *E. coli* are responsible for a wide range of infections, including pneumonia, septicemia, and urinary tract infections, occurring in both community and healthcare settings (7). The widespread dissemination of CPE is largely driven by the horizontal transfer of antibiotic resistance genes through mobile genetic elements such as plasmids and transposons (8). Monitoring the global dissemination of these mobile elements and their association with carbapenemase genes in *E. coli* and *K. pneumoniae* clones remains a critical public health priority, essential for developing effective containment, management, and prevention strategies (9).

Despite the numerous reports of CPE in Peru over the past twelve years, comprehensive genomic data from longitudinal and multicenter studies remain lacking—data that are essential to contextualize local epidemiology within regional and global frameworks. Therefore, the aim of this study was to characterize the phenotypic and genomic features of *Escherichia coli* and *Klebsiella pneumoniae* clinical isolates producing carbapenemases, collected between 2020 and 2022 from four healthcare institutions in Lima, Peru.

## MATERIALS AND METHODS

### Clinical isolates

Between 2020 and 2022, a total of 61 *E. coli* and 259 *K. pneumoniae* non-redundant, unique suspected CPE isolates were selected based on the presence of an ertapenem inhibition zone ≤ 22 mm. Clinical isolates were obtained from urine and blood samples collected from both inpatients and outpatients and processed at the microbiology laboratories of four healthcare institutions in Lima, Peru: Hospital de Emergencias Ate Vitarte, Hospital Nacional Guillermo Almenara Irigoyen, Hospital Nacional Hipólito Unanue, and Laboratorio Clínico ROE (Table 1). All isolates were subsequently sent to the Universidad de Piura (UDEP) for phenotypic and genomic characterization.

**Table 1.** Distribution of clinical isolates by bacterial species, healthcare institution, and sample type.

| Bacterial species | Health institutions * |  |  |  |  |  |  |  |
| --- | --- | --- | --- | --- | --- | --- | --- | --- |
|  | HEA |  | HHU |  | HGA |  | ROE |  |
|  | Urine | Blood | Urine | Blood | Urine | Blood | Urine | Blood |
| *E. coli* | 1 | 0 | 1 | 0 | 20 | 0 | 39 | 0 |
| *K. pneumoniae* | 9 | 3 | 35 | 14 | 64 | 54 | 78 | 2 |

\*Hospital de Emergencias Ate-Vitarte (HEA), Hospital Nacional Hipólito Unanue (HHU), Hospital Nacional Guillermo Almenara Irigoyen (HGA), y Laboratorio Clínico Roe (ROE)

### Carbapenemase screening and identification

The phenotypic screening of carbapenemase production was done with the Blue carba Test as previously described (10). Inhibition assays using EDTA, phenylboronic acid, and avibactam were performed to differentiate among Ambler classes of carbapenemasas (11).

### Antimicrobial susceptibility testing and molecular detection of carbapenemase genes

Resistance phenotypes were determined using the disk diffusion method and interpreted according to the clinical breakpoints established by the *Clinical and Laboratory Standards Institute (CLSI) M100* guidelines (12). Colistin susceptibility was assessed by the COL-spot test, as previously described (13). Intermediate resistance results were interpreted and reported as resistant, and isolates were classified as multidrug-resistant (MDR), extensively drug-resistant (XDR), or pandrug-resistant (PDR) according to the criteria proposed by Magiorakos *et al.* (14). For *K. pneumoniae*, fosfomycin susceptibility was interpreted using the cutoff values established for *E. coli* in the CLSI M100 guideline (12).

The minimum inhibitory concentration (MIC) was determined in 100 selected isolates against aztreonam/avibactam, ceftazidime/avibactam (agar dilution), and cefiderocol (broth microdilution), following CLSI M100 recommendations (12). Isolate selection was based on carbapenemase type, maintaining proportional representation (*bla*_KPC_ = 30, *bla*_NDM_ = 30, *bla*_OXA-48-like_ = 30, *bla*_NDM_ + *bla*_KPC_ = 9 y *bla*_NDM_ + *bla*_OXA-48-like_ = 1). Carbapenemase genes were identified by polymerase chain reaction (PCR) amplification as previously described (15).

### Whole-Genome Sequencing (WGS)

Isolates considered of greatest epidemiological or microbiological interest—particularly those harboring multiple resistance determinants—were selected for whole-genome sequencing (WGS). A total of 23 *E. coli* and 60 *K. pneumoniae* isolates underwent short-read sequencing using the Illumina NextSeq platform (paired-end, 2 × 250 bp).

From these, five *E. coli* and three *K. pneumoniae* isolates were additionally selected for long-read sequencing using the MinION platform (Oxford Nanopore Technologies, UK). Genomic libraries were prepared with the Nextera XT DNA Library Preparation Kit (Illumina, UK) for short reads and the Ligation Sequencing Kit with Library Loading Beads (Oxford Nanopore Technologies, UK) for long reads, following the manufacturer’s protocols.

### Bioinformatics analysis

Raw sequence reads were assessed for quality using FastQC v0.11.9 (16). Low-quality reads (Phred score < 30) and adapter sequences were trimmed using Trimmomatic v0.39 (17). *De novo* genome assemblies were generated with Unicycler v0.4.8 (18), and assembly quality was evaluated using QUAST v5.0.2 (19). Genome annotation was performed with Prokka v1.14.6 (20) and subsequently curated manually.

The resulting assemblies were analyzed using tools from the Center for Genomic Epidemiology (http://www.genomicepidemiology.org/) and other publicly available platforms. Specifically, multilocus sequence types (MLSTs) were identified using MLSTFinder, antimicrobial resistance genes with ResFinder, plasmid replicons with PlasmidFinder and pMLSTFinder, insertion sequences and transposons with ISFinder (https://isfinder.biotoul.fr/), and virulence factors using Kleborate for *K. pneumoniae* and VirulenceFinder for *E. coli*.

Phylogenetic grouping of *E. coli* was determined with ClermonTyping (http://clermontyping.iame-research.center/). Serotype prediction was conducted using SeroTypeFinder for flagellar (H) and lipopolysaccharide (O) loci in *E. coli*, and Kaptive for capsule (K) and lipopolysaccharide (O) loci in *K. pneumoniae*. Prophage regions were identified using PHASTER (https://phaster.ca/), and genomic context analyses were supported by Pathogenwatch (https://pathogen.watch/).

Single-nucleotide polymorphism (SNP)–based phylogenomic relationships were inferred using SNP-sites v2.5.1 to extract SNP positions, followed by maximum-likelihood tree construction with IQ-TREE v1.5.5.3, applying the best-fit substitution model and 1,000 bootstrap replicates for node support. Tree was visualized with Microreact (https://microreact.org).

### Data Availability

Whole genome sequencing data have been deposited in the NCBI Sequence Read Archive (SRA) under BioProject accession numbers PRJNA1427713 (*Klebsiella pneumoniae*) and PRJNA1265301 (*Escherichia coli*).

## RESULTS

### Phenotypic screening and antimicrobial susceptibility profiles

Phenotypic screening for carbapenemase production identified 222 metallo-β-lactamase (MBL) producers (*E. coli*, n = 41; *K. pneumoniae*, n = 181), 54 class A serine carbapenemase producers (*E. coli*, n = 1; *K. pneumoniae*, n = 53), and 33 class D serine carbapenemase producers (*E. coli*, n = 19; *K. pneumoniae*, n = 14). Notably, 12 isolates were positive for both serine and metallo-carbapenemase activity according to phenotypic assays.

The antimicrobial susceptibility profiles of the 61 *E. coli* isolates are summarized in Figure 1. Carbapenem resistance was variable, with some isolates remaining susceptible—likely associated with the presence of class D OXA-48-like enzymes, which often confer lower levels of resistance. High resistance rates were observed for ciprofloxacin (61/61, 100%), trimethoprim/sulfamethoxazole (51/61, 83.6%), amikacin (46/61, 75.4%), and ceftazidime/avibactam (41/61, 67.2%), alongside widespread resistance to cephalosporins and monobactams. Conversely, lower resistance rates were recorded for fosfomycin (4/61, 6.5%) and colistin (5/61, 8.2%).

**Figure 1.**
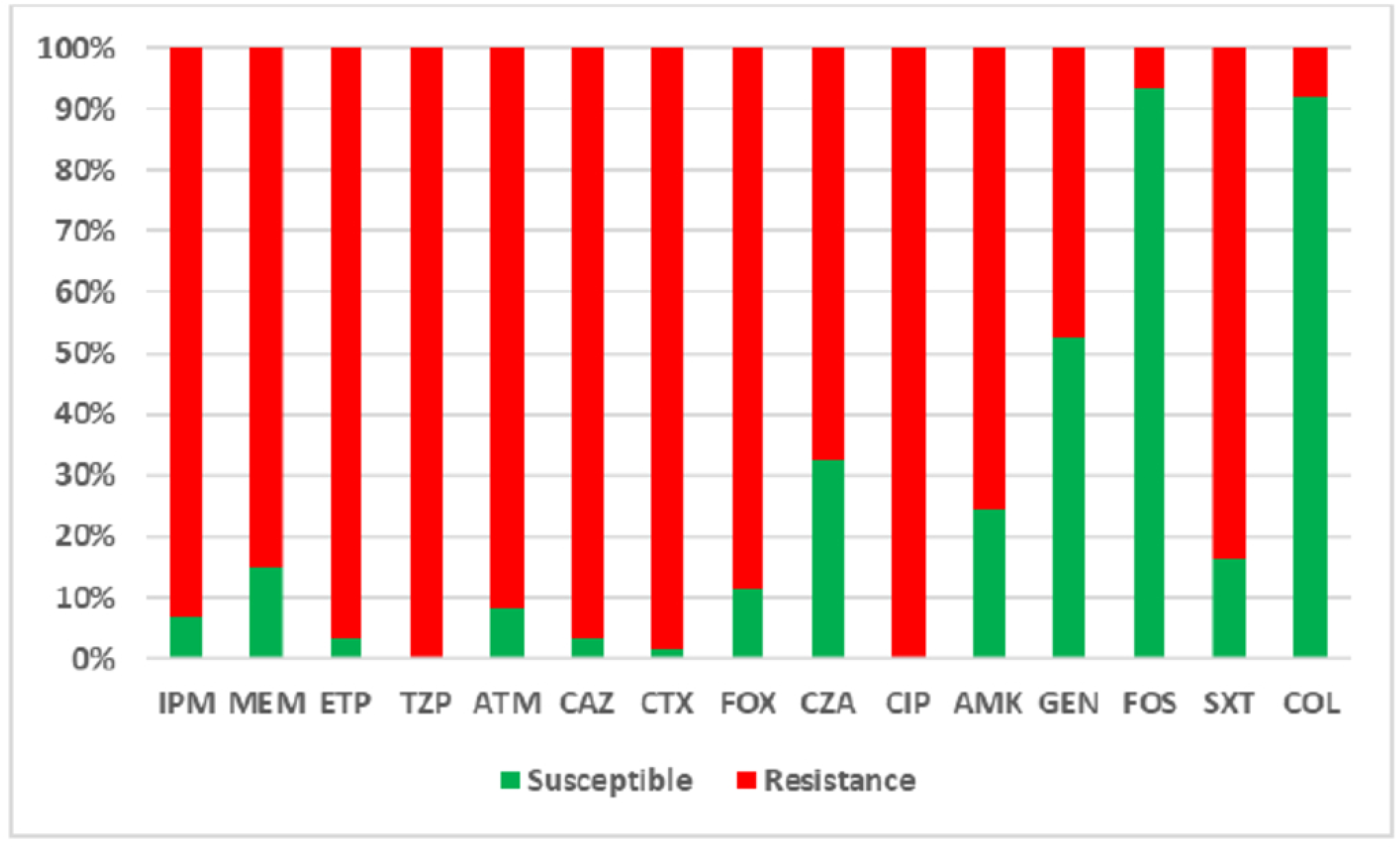
Distribution of antimicrobial resistance among carbapenemase-producing *Escherichia coli* clinical isolates.

Regarding the antibiotic susceptibility profile of the 259 *Klebsiella pneumoniae* isolates (Figure 2), in addition to widespread resistance to carbapenems, nearly all isolates were resistant to cephalosporins, monobactams, and fluoroquinolones. High resistance rates were also observed for trimethoprim/sulfamethoxazole (212/259, 81.9%), gentamicin (231/259, 89.2%), amikacin (170/259, 65.6%), and ceftazidime/avibactam (192/259, 74.1%). Conversely, lower resistance levels were detected for fosfomycin (38/259, 14.7%) and colistin (76/259, 29.3%).

**Figure 2.**
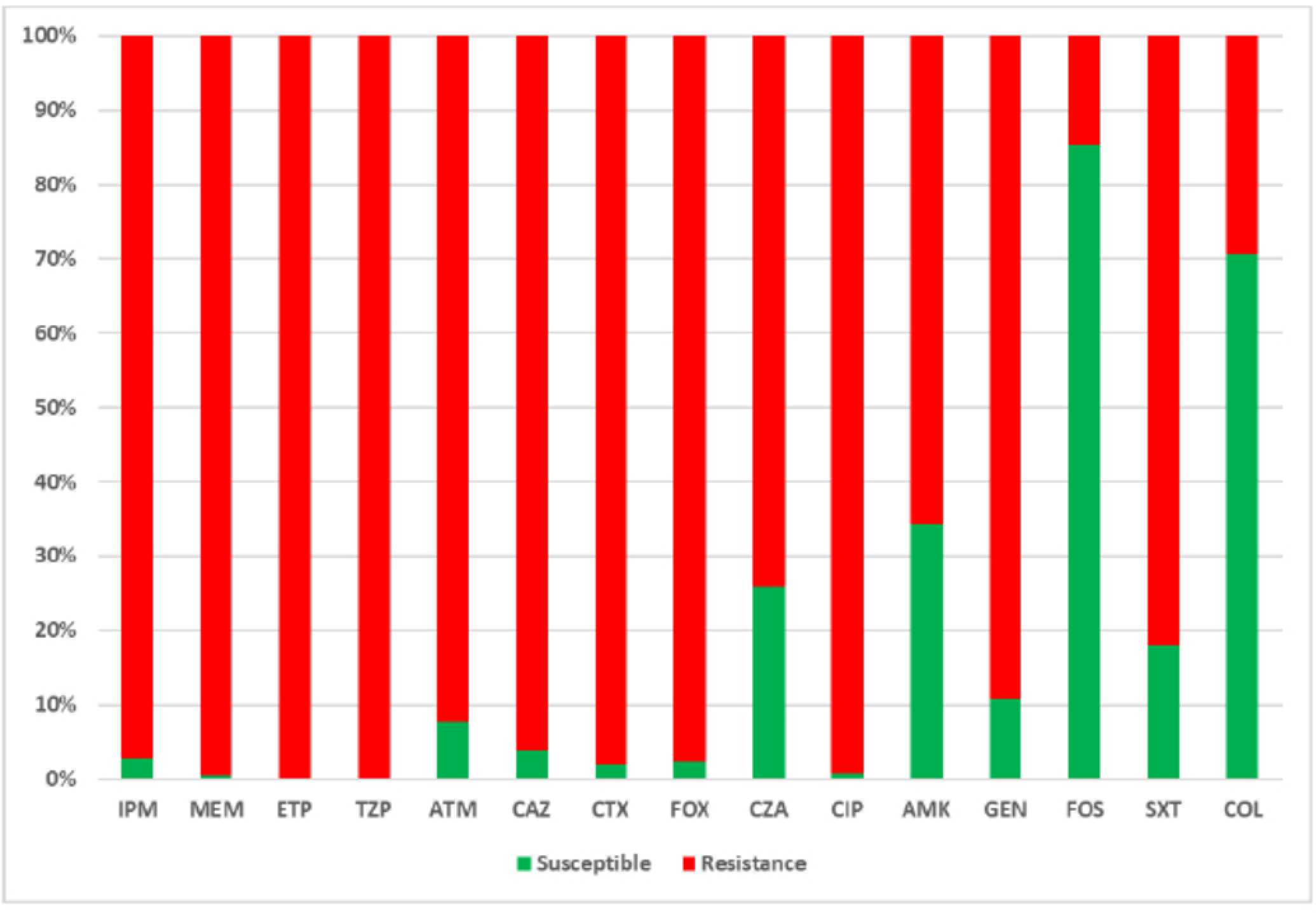
Distribution of antimicrobial resistance among carbapenemase-producing *Klebsiella pneumoniae* clinical isolates. 1PM: imipenem, MEM: meropenem; ETP: ertapenem: TZP: piperaclllin/tazobactam; ATM: aztreonam; *CAZ:.* ceftazidime; en<: cefotaxime; FOX: cefoxitin; CZA ceflazidime/avibactam; CIP: ciprofloxacin; AMK: amikacin, GEN: gentamicin; FOS: fosfomycin; SXT: sulfamethoxazole/trimethoprim; COL: colistin.

It is noteworthy that only serine carbapenemase-producing isolates remained susceptible to ceftazidime/avibactam, whereas—as expected—isolates producing exclusively metallo-β-lactamases (MBLs) or OXA-48-like enzymes, which lacked extended-spectrum β-lactamases (ESBLs), retained susceptibility to aztreonam and third-generation cephalosporins, respectively.

A total of 17 isolates (1 *E. coli* and 16 *K. pneumoniae*) were resistant to all antimicrobials tested and were thus classified as pandrug-resistant (PDR). One hundred one isolates (10 *E. coli* and 91 *K. pneumoniae*) were categorized as extensively drug-resistant (XDR), while the remaining carbapenemase-producing *Enterobacterales* were classified as multidrug-resistant (MDR) according to international criteria.

### Minimum Inhibitory Concentration (MIC) for selected β-lactams

The MIC results for cefiderocol, ceftazidime/avibactam, and aztreonam/avibactam among the 100 selected isolates are summarized in Table 2. Notably, all isolates were susceptible to aztreonam/avibactam and cefiderocol, with no resistance detected. In contrast, 40% of the isolates exhibited resistance to ceftazidime/avibactam, a finding that correlated with the presence of the *bla*_NDM_ gene..

**Table 2.** Susceptibility profile of selected isolates of *E. coli* and *K. pneumoniae* against new β-lactams antimicrobials.

| **Carbapenemase Class** | **Number of Isolates (%)** | **Ceftazidime/Avibactam (%)** | **Aztreonam/Avibactam (%)** | **Cefiderocol (%)** |
| --- | --- | --- | --- | --- |
| *Total* | 91 (100) | 59.3 | 100 | 100 |
| *A* | 28 (31) | 100 | 100 | 100 |
| *B* | 27 (30) | 0 | 100 | 100 |
| *D* | 26 (28) | 100 | 100 | 100 |
| *A + B* | 9 (10) | 0 | 100 | 100 |
| *B + D* | 1 (1) | 0 | 100 | 100 |

\* The outcome could not be determined in 9 isolates.

### Genetic markers of carbapenem resistance

Detection of carbapenemase-encoding genes by PCR allowed the recognition of 69.0% isolates carrying *bla*_NDM_, (40 *E. coli* and 181 *K. pneumoniae*), 16.9% isolates carrying *bla*_KPC_ (1 *E. coli* and 53 *K. pneumoniae*) and 4.6% isolates carrying *bla*_OXA-48-like_ (19 *E. coli* and 14 *K. pneumoniae*). Furthermore, 11/320 (3.4%) *K. pneumoniae* isolates tested positive for *bla*_NDM_ and *bla*_KPC_ genes and one *E. coli* isolate was co-carrying *bla*_NDM_ and *bla*_OXA-48-like_, corresponding with the phenotypic results, excepting for the isolate of *E. coli* which was phenotypically detected only as MBL (Table 3).

**Table 3.** Distribution of carbapenemase genes among *Escherichia coli* and *Klebsiella pneumoniae* clinical isolates.

|  | ***bla*<sub>KPC</sub>** | ***bla*<sub>NDM</sub>** | ***bla*<sub>OXA-48-like</sub>** | ***bla*<sub>NDM</sub> + *bla*<sub>KPC</sub>** | ***bla*<sub>NDM</sub> + *bla*<sub>OXA-48-like</sub>** | **Total** |
| --- | --- | --- | --- | --- | --- | --- |
| *E. coli* | 1 | 40 | 19 | 0 | 1 | 61 |
| *K. pneumoniae* | 53 | 181 | 14 | 11 | 0 | 259 |
| *Total* | 54<br>(16.9%) | 221<br>(69.1%) | 33<br>(10.3%) | 11<br>(3.4%) | 1<br>(0.3%) | 320<br>(100.0 %) |

### Analysis of Whole Genome Sequencing

WGS identified that *E. coli* (n=23) isolates harbored an average of 12 antimicrobial resistance (AMR) genes (SD = 4.4). The genes *bla*_OXA-1_, *aac(6′)-lb-cr*, *sul-1*, *sul-2*, and *catB3* were present in over 78% of *E. coli* isolates. The genes responsible for AMR among CPE isolates are shown in Figure 3. Regarding the genes encoding carbapenemases, two allelic variants of *bla*_NDM_ were detected: *bla*_NDM-5_ (10/23, 43.5%) and *bla*_NDM-1_ (8/23, 39.1%); *bla*_OXA-48_ (4/23, 17.3%) was also detected. One (4.3%) isolate tested positive for *bla*_NDM-1_ and *bla*_OXA-48_ genes.

**Figure 3.**
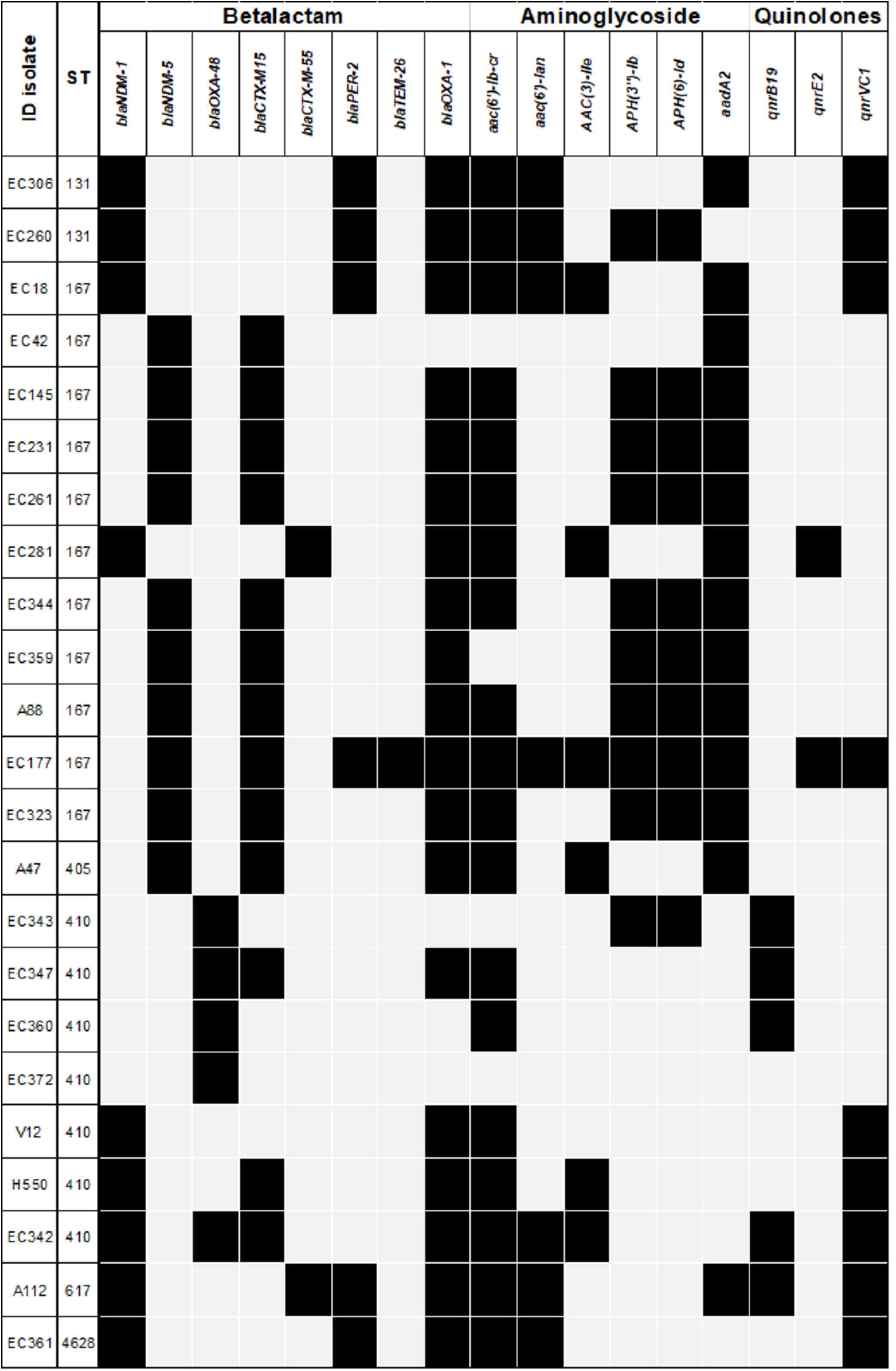
Resistance genes of carbapenemase-producing *Escherichia coli.* The mam antimicrobial resistance **(AMR)** genes are summarized by bacterial isolate and the most important antimicrobial families.

WGS was performed on 60 *K. pneumoniae* isolates, which harbored an average of 13 (SD = 2.8) AMR genes. The *bla*_OXA-1_ (57/60, 95%), *aac(6′)-lb-cr* (56/60, 93.3%) and *sul-1* (56/60, 93.3%) genes were detected in nearly all isolates. Several AMR genes responsible for resistance to non-carbapenem β-lactams, including AmpC β-lactamases, extended-spectrum β-lactamases (ESBLs), aminoglycosides, fluoroquinolones, and other antimicrobial classes, were also detected among CPE isolates and are shown in Figure 4. Regarding carbapenemase-encoding genes, *bla*_NDM-1_ (53/60, 88.3%) was detected in almost all isolates, whereas bla_KPC-2_ (15/60, 25%) and *bla*_OXA-181_ (1/60, 1.6%) also were detected. Nine (15%) isolates tested positive for both *bla*_NDM-1_ and *bla*_KPC-2_ genes.

**Figure 4A.**
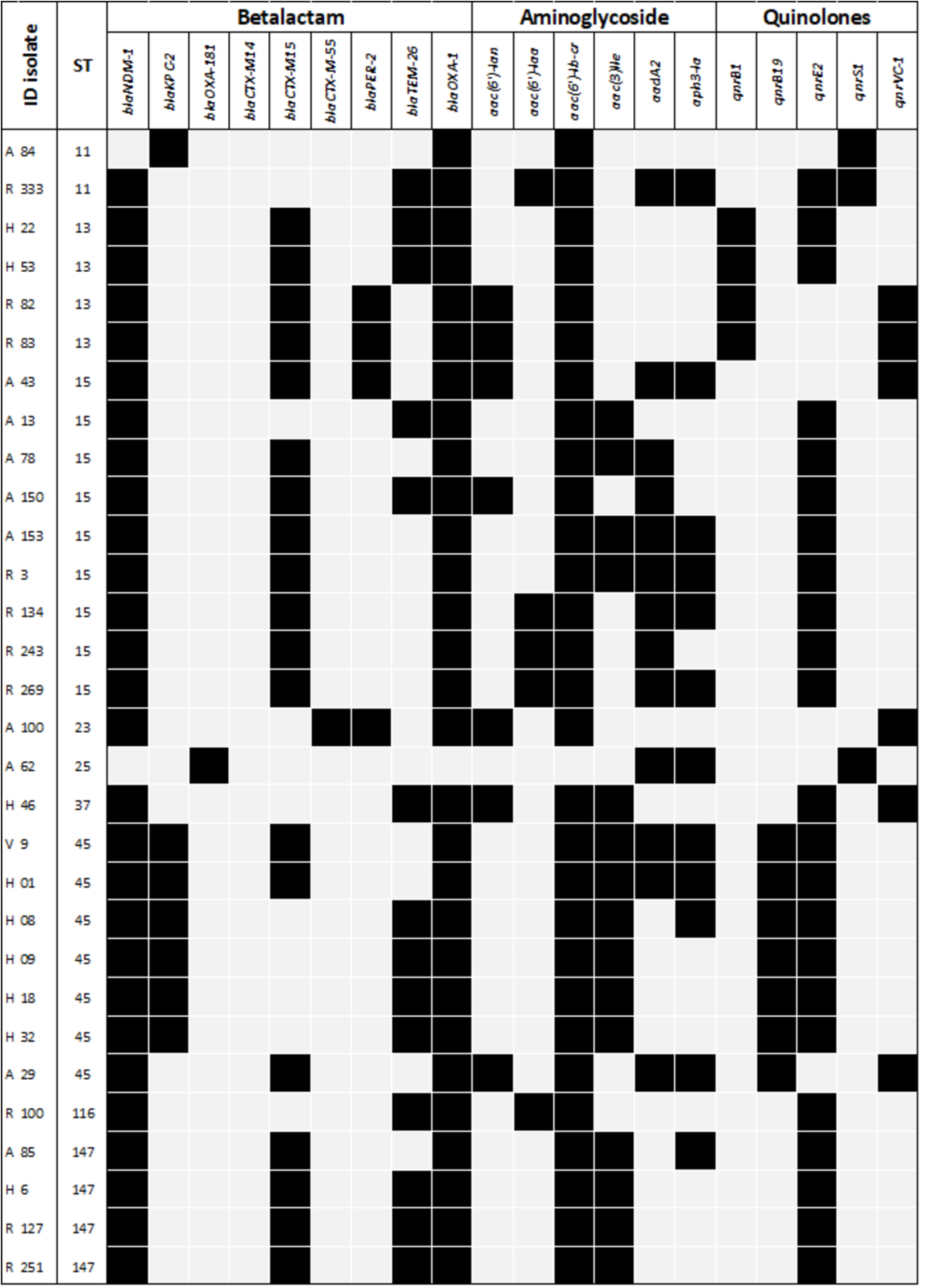
Resistance genes of carbapenemase-producing Klebsiella pneumoniae. The main antimicrobial resistance **(AMR)** genes are summarized by bacterial isolate and the most important antimicrobial tamilies

**Figure 4B.**
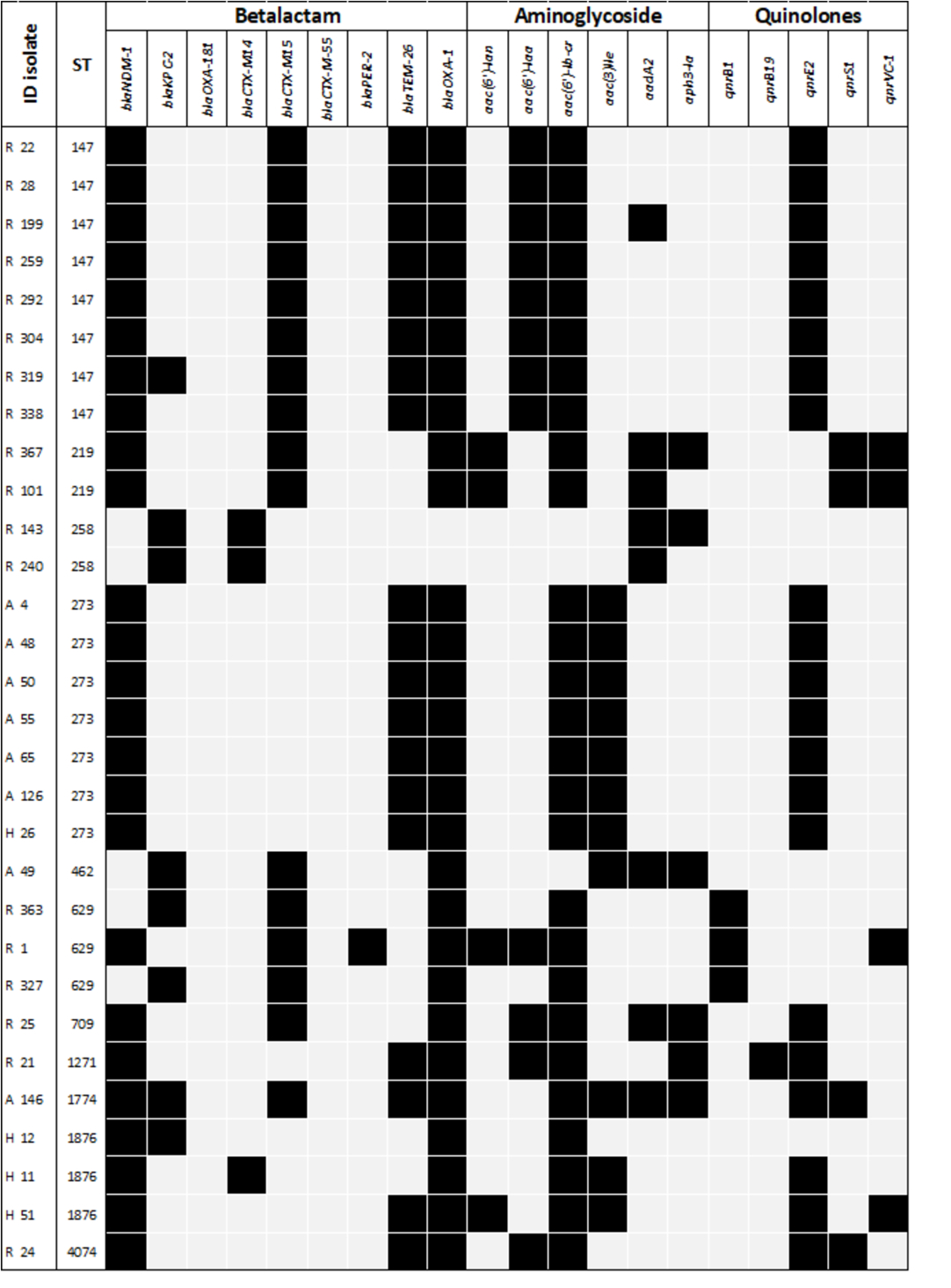
Resistance genes of carbapenemase-producing Klebsiella pneumoniae. The main antimicrobial resistance (AMR) genes are summarized by bacterial isolate and the most, important antimicrobial families.

### Bacterial population structure and linkage to specific carbapenemase alleles

Six diverse genetic backgrounds were identified among the *E. coli* isolates (n = 23). The most frequent sequence types were ST167 (n = 11), ST410 (n=7) and ST131 (n = 2), each represented by more than one isolate. All eleven ST167 isolates carried *bla*_NDM_, specifically *bla*_NDM-5_ (n = 9) and *bla*_NDM-1_ (n = 2). Among the seven ST410 strains, four harbored *bla*_OXA-48_, two *bla*_NDM-1_ and one co-harbored both *bla*_OXA-48_ and *bla*_NDM-1_. Both ST131 isolates carried *bla*_NDM-1_. The remaining isolates exhibited greater genetic diversity, carrying different *bla*_NDM_ variants: *bla*_NDM-1_ was identified in ST617 and ST4628 and *bla*_NDM-5_ was present in ST405. Phylogenetic analysis revealed a close genetic relationship among ST167 isolates, with a divergence of 6 to 21 SNPs, while ST410 isolates exhibited a divergence of only 2 to 3 SNPs.

MLST showed that *K. pneumoniae* was dominated by NDM-producing clonal group (CG) 147, more specifically ST147 (n = 12) and its single locus variants ST273 (n = 7). The CG147 cluster comprised 19 isolates, all NDM-producers, and one of the ST147 isolates was a co-producer of NDM and KPC. Among the *K. pneumoniae* isolates MLST identified other clusters represented by more than one isolate: one representing CG15 and including ST15 (n=9) and ST709 (n=1), all *bla*_NDM-1_-harboring; other representing CG45 and including ST45 (n=7) with co-harboring *bla*_NDM-1_ and *bla*_KPC-2_ (n = 6) and one only with *bla*_NDM-1_; other representing CG258 and including ST258 with *bla*_KPC-2_ (n = 2) and ST11 with *bla*_KPC-2_ (n=1) and *bla*_NDM-1_ (n = 1); other representing CG13 and including ST13 with *bla*_NDM-1_ (n = 4); other representing CG629 and including ST629 with *bla*_KPC-2_ (n=2) and *bla*_NDM-1_ (n = 1); other representing ST1876 (n=3) with co-harboring *bla*_NDM-1_ and *bla*_KPC-2_ (n = 1) and with *bla*_NDM-1_ (n=2); and other representing CG107 and including ST219 (n=2) with *bla*_NDM-1_. The remaining *K. pneumoniae* isolates represented genetically diverse single strains harboring *bla*_NDM-1_ (ST23, ST37, ST116, 1271 and ST4074), *bla*_NDM-1_ + *bla*_KPC-2_ (ST1774), *bla*_KPC-2_ (ST462), and *bla*_OXA-181_ (ST25). Among the most frequent STs, we observed that most *K. pneumoniae* ST147 genomes diverged by 12 to 104 SNPs. However, two isolates were more distant, with 500–2831 SNPs. ST15 isolates diverged by 12 to 45 SNPs, although three isolates were more distant, with 842–1704 SNPs. ST45 isolates diverged by 11–427 SNPs, and most ST273 isolates diverged by 2–20 SNPs. However, one isolate showed a distance greater than 2800 SNPs.

### Replicon Typing

PlasmidFinder showed that in *E. coli* isolates the most frequent replicon were IncFII, IncFIA, and IncFIB, representing 87% each, followed by IncC with 39.1%, Col(pHAD28) with 26% (related to the *qnrB19* gene, which generates low-level quinolone resistance) and IncY with 17.4%. Other replicons were also detected. However, it was in a proportion less than 15%, such as P0111, IncFIB/HI1B, IncI-1, Col (BS512), IncX1, IncX4, IncFIC, and IncR. Virtually all isolates (n = 22/23; 95.6%) presented a multireplicon state, with three or more different Inc groups.

Regarding STs and carbapenemase-related replicons, we observed that ST167 (n=11) was associated mostly with hybrid plasmid IncFIA/FIB/FII *bla*_NDM-5_-harboring (n=9), and other two associated with IncFIB/HI1B and IncC2 *bla*_NDM-1_-harboring; three ST410 isolates was associated with IncC2 *bla*_NDM-1_-harboring. Highlight, that *bla*_OXA-48_ was not located on a plasmid but on the chromosome of ST410 isolates.

Nineteen diverse Inc groups were identified in *K. pneumoniae*, IncFIB/HI1B and IncFIB(K) were the most common Inc group with 75% each, followed by IncFII(K), IncC2, colRNAl, col(pHAD28), IncFll, IncFlB and IncU/IncX3 found in 48.3%, 31.7%, 23.3%, 21.7%, 20.0%, 18.3%, and 16.7% of isolates, respectively. Other replicons were also detected. However, it was in a proportion less than 15%, such as IncFlA, col4401, IncM, IncN, IncX, IncR, IncX3, and IncHI2. The majority of isolates (n = 54/60; 90%) were characterized by a multireplicon status carrying three or more different Inc groups.

Regarding STs and carbapenemase-related replicons, we observed that ST147 (n = 12) *bla*_NDM-1_-harboring, were associated with IncFIB/HI1B, and one of them (co-producing carbapenemases KPC and NDM) also was associate with IncU/IncX3; ST15 *bla*_NDM-1_-harboring was associated mostly with IncFIB/HI1B (n = 8) and with IncC2 (n = 1); ST273 *bla*_NDM-1_-harboring was associated with IncFIB/HI1B. Finally, ST45 *bla*_NDM-1_ and *bla*_KPC-2_-harboring was associated mostly with IncFIB/HI1B and IncU/IncX3, respectively (n = 6) and one isolate that only *bla*_NDM-1_-harboring was associated with IncC2.

### Serotype and Phylogenetic group of *Escherichia coli*

The lipopolysaccharide (O) and flagellar (H) surface antigens of *E. coli* are targets for serotyping that have traditionally been used to identify pathogenic lineages. These surface antigens are important for the survival of *E. coli*. Other strategy is phylogroup classification, allowing the identification of seven phylogroups designated as A, B1, B2 C, D, E, and F. The use of phylogroup classification has been employed in the study of ecological niches and lifestyles in bacterial pathogens. Eleven ST167 isolates showed a O101:H10 (10) serotype and O101:H4 (1) serotypes, only one O101:H10 serotype presented the fimbria variant fimH54, the others were negative, all belonged to phylogroup A. The ST410 belonged to the O8:H21 serotype with fimH24 and phylogroup C, the ST131 belonged to the O25:H4 with fimH30 and phylogroup B2, the ST405 belonged to O102:H6 with fimH27 and phylogroup D, the ST617 belonged to H9 within fimH and phylogroup A, and the ST4628 belonged to H11 within fimH and phylogroup A.

### K and O loci of Klebsiella pneumoniae

The K-Locus capsular polysaccharide (CPS) and the O-Locus lipopolysaccharide (LPS) are important determinants of virulence and bacterial interaction with the immune system. We identified 21 different K loci and 8 O loci, and these K/O loci provided 32 different combinations in our *K. pneumoniae* isolates. KL64 (21.6%), KL74 (11.6%), KL62 (11.6%), and KL48 (10%) were the most prevalent K loci. Among the predominant STs, we observed the following K+O loci: KL64+O1/O2v1 (21.6%, 13/60) associated with ST147 (n = 11) and ST15 (n = 2); KL74+OL104 (11.6%, 7/60) associated with ST243; KL62+O1/O2v1 (11.6%, 7/60) associated with ST45; and KL48+O1/O2v2 (6.6%, 4/60) associated with ST15.

### Virulence factors

All *E. coli* isolates presented virulence genes, harboring an average of 17 (SD = 7) virulence genes. A total of 53 virulence genes were identified. The following genes were found in 80% of the isolates: the fimbrial genes *yeh*A/C/D (responsible for adhesion to some abiotic surfaces), *terC* (a tellurium ion resistance protein), *fdeC* and *csgA* (adhesins), *hha* (which modulates the expression of biofilm formation), and *hlyE* (hemolysin). In addition, we identified other important virulence genes, including *fimH*, associated with type 1 fimbriae, in all ST410 and ST131 isolates, and one ST167 isolate; *iucC*, related to aerobactin synthetase, *iutA*, encoding the aerobactin ferric receptor, and *sitA*, associated with iron acquisition, in all isolates of ST410 and ST131, in one ST167 and another ST617. Highlight, that the two ST131 isolates presented more than 30 virulence genes.

In addition to the capsular polysaccharide (K antigen) and LPS (O antigen) biosynthesis loci, a set of virulence factors was described, which were present in 45% (27/60) of *K. pneumoniae* isolates.

The most prevalent virulence factor was the yersiniabactin gene cluster (85.2%; 24/27). The majority of yersiniabactin-positive (*ybt_+_*) isolates, 59.3% (16/27), were spread via ICE*Kp4* related to *ybt1-* lineage, corresponding mostly to ST45 (n=7) and ST15 (n=6) isolates. Second, 14.8% (4/27) *ybt_+_* isolates revealed an ICE*Kp3* (*ybt* lineage 9), associated mostly with ST11 isolates (n = 2). We also identified 7.4% (2/27) of unassigned *ybt* lineage (*ybt0*), associated with ST13 isolates. In addition, we found aerobactin *iuc5* associated with ST147 (n=4). Highlight, the presence of an isolate belonging to ST23, which usually possess several virulence factors, this showed ICE*Kp1-* related to *ybt1* lineage, colibactin *clb2*, aerobactin *iuc1*, salmochelin *iro1* and genes upregulators of capsule expression rmp1/A2.

### Genetic context of carbapenemase genes

We analyzed the genetic environment of the sequenced and assembled isolates by hybrid assembly.

*bla*_NDM-1_ in plasmid IncFIB/IncHI1B.-The plasmid pKp319ndm *bla*_NDM-1_-harboring, was located in very large IncFIB/HI1B hybrid plasmid, which size was ∼359.7 kb. A common genetic environment around *bla*_NDM-1_ (IRR of IS*Aba125*-*bla*_NDM-1_-*ble*_MBL_-*trp*F-*dsb*C-*dct*-*gro*ES) was identified; this region seems to be flanked by two IS*Kox2-*like insertion sequences. In addition, other rearrangements of resistance determinants and mobile elements were described downstream [*Tn2*-like-*bla*_TEM-26_-*aac*(*3*)*-IIe*-IS*Kpn11*-IS*Kpn12*-IS*1R*-*qnr*E2-IS*Ecp1*] and upstream [IS*1R*-*cat*A1-*Tn3*-like-IS*26*-*intI1*-*aac(6’)-lb-cr-bla*_OXA-1_-*cat*B3-*arr-3*-*qac*E-*sul1*-IS*1326*] of the previous genetic environment.

*bla*_NDM-1_ in plasmid IncC2.-The plasmid pEc306ndm belonged to IncC type 2 incompatibility group. It has a size of ∼203.2 kb. The *bla*_NDM-1_ gene is flanked upstream by an IRR of ISAba125 and downstream by the bleomycin resistance gene *ble*_MBL_, forming, together with *trp*F-*dsb*C-*dct*-*Δgro*ES, a common genetic environment around *bla*_NDM-1_, (IRR of IS*Aba125*-*bla*_NDM-1_-*ble*_MBL_-*trp*F-*dsb*C-*dct*-*gro*ES) that is similar to that described previously. Furthermore, it was observed; upstream, a zone delimited by two copies of IS*Kox2*-like elements carrying the *bla*_PER-2_ gene previously described environment around this gene (IS*Pa12*-*bla*_PER-2_-*gst*-like-*abct*), followed by a region where a class 1 integron [*aac(6’)-Ib-cr*-*bla*_OXA-1_-*cat*B3-*arr-3*-*sul1*] was present. Finally, downstream, the *mer* operon (*mer*RTPCA) was present, which confers resistance to mercury compounds through the expression of the MerA protein.

*bla*_NDM-5_ in plasmid IncFIA/IncFIB/IncFII.-The *bla*_NDM-5_ gene, from a hybrid genomic assembly, was identified in a mosaic plasmid (pEc231ndm), which presented a size of ∼140.3 kb, with three origins of replication (IncFIA/IncFIB/IncFII). The *bla*_NDM-5_ gene is flanked upstream by an IRR of IS*Aba125* and downstream by the bleomycin resistance gene *ble*_MBL_, forming, together with *trpF-dsbC*, a common genetic environment, that is similar to that described previously to *bla*_NDM-1_ (IRR of IS*Aba125*-*bla*_NDM-5_-*ble*_MBL_-*trpF-dsbD*). Furthermore, it was observed; upstream, a class 1 integron (*dfrA12-aadA2-qacC-sul1*). Finally, downstream, a zone delimited by two copies of IS*26*-like elements carrying *bla*_CTX-M-15_*-catB3-bla*_OXA-1_*-aac(6’)-Ib-cr* genes.

*bla*_KPC-2_ in plasmid IncX3/IncU.-The plasmid pKp319kpc *bla*_KPC-2_-harboring, was located in IncX3/IncU hybrid plasmid, which size was ∼46.6 kb. The genetic context of *bla*_KPC-2_ gene was harbored by a non-Tn4401 element (NTE_KPC_) classified as NTE_KPC_-_Ic_, presenting a partial IS*Kpn6* with the associated left inverted repeat and a Tn3 resolvase gene (*tnp*R) downstream and upstream, respectively, of *bla*_KPC-2_.

*bla*_OXA-48_ in chromosome.-From the hybrid assembly in EC347, it was located that the *bla*_OXA-48_ gene was fragment was located in the chromosome, flanked by two copies of IS*1R*. Furthermore, it was observed; upstream, a copper resistance determinant, which contains six genes, *copABCDRS*, arranged in two operons, *copABCD* and *copRS*; and downstream, a *NikABCDE* system, belongs to the ABC transporter family, composed of the periplasmic binding protein NikA, two integral membrane components (NikB and -C), and two ATPases (NikD and -E).

## DISCUSSION

This study aimed to evaluate the epidemiology of carbapenemase-producing *Escherichia coli* and *Klebsiella pneumoniae* recovered over a three-year period (2020–2022) from four healthcare institutions in Lima, Peru, revealing the emergence and multiclonal dissemination of these pathogens. The spread of CPE is now widely established across Latin America and is frequently associated with the co-occurrence of resistance mechanisms to multiple antimicrobial classes, which significantly complicates clinical management of infections (21).

All carbapenemase-producing isolates in this study displayed a MDR phenotype, underscoring the potential limitations in available therapeutic options and highlighting the urgent need for a coordinated global response to address this emerging threat. Among the agents tested, ciprofloxacin, trimethoprim/sulfamethoxazole, and amikacin exhibited the lowest activity, with resistance rates exceeding 75%, and reaching 100% for ciprofloxacin. These findings confirm the limited empirical utility of fluoroquinolones and trimethoprim/sulfamethoxazole for the management of urinary tract infections caused by CPE, in line with the Infectious Diseases Society of America (IDSA) recommendations, which advise against the empirical use of these agents in settings where local resistance rates surpass 20% (22).

Drugs commonly employed in combination therapies for CPE infections showed variable activity against contemporary isolates. In our cohort, fosfomycin exhibited low resistance rates (<15%; 14.3% in *K. pneumoniae* and 6.6% in *E. coli*), comparable to data reported from Argentina (11.4%) (23), Mexico (10.9%), and China (10%) (24, 25). However, these results contrast with previous reports from Peruvian hospitals, where fosfomycin resistance reached 27.8% (26), and with studies from Southeast Asia and North Africa documenting prevalence up to 36.4% in Vietnam and 38.5% in Egypt (27, 28).

Regarding colistin, resistance rates reached 29.3% in *K. pneumoniae* and 8.2% in *E. coli*. Although colistin remains a last-resort option, its efficacy has been increasingly compromised by the rising prevalence of resistant strains across Latin America (29). A recent multicenter study in Peru analyzing 317 clinical isolates from 12 regions reported lower colistin resistance rates—7.5% in *E. coli* and 11.9% in *K. pneumoniae* (30)—values below those observed in the present study. This difference may be explained by our focus on carbapenemase-producing isolates, which have been reported to exhibit colistin resistance rates as high as 31% (31).

Given the limited therapeutic options available for severe infections caused by CPE, combination regimens including colistin and fosfomycin may provide valuable alternatives against these multidrug-resistant clinical isolates (32).

The high rate of amikacin resistance observed in this study is particularly concerning, given that this aminoglycoside has traditionally been used in Peru as a reserve agent against multidrug-resistant (MDR) Gram-negative bacilli and as an alternative therapeutic option in settings where access to newer antimicrobial agents remains limited. In *E. coli*, resistance to amikacin was higher than to gentamicin (75.4% vs 47.5%), whereas in *K. pneumoniae*, the opposite pattern was observed, with gentamicin showing higher resistance than amikacin (89.2% vs 65.6%). This discrepancy may reflect the presence of distinct aminoglycoside resistance genes conferring selective resistance to each compound (32).

Recent studies recommend the primary use of novel β-lactam/β-lactamase inhibitor combinations such as ceftazidime/avibactam (CZA) for infections caused by serine carbapenemase producers, and aztreonam/avibactam (AZA) or cefiderocol for those mediated by metallo-β-lactamases (MBLs) (32). In agreement with these recommendations, our study demonstrated complete susceptibility (100%) to CZA among serine carbapenemase producers, consistent with data from recent global surveillance programs that included isolates from Latin America (33, 34). However, although no resistance to CZA was detected among serine carbapenemase producers in our collection, reports from Peru have described CZA-resistant *Klebsiella pneumoniae* associated with KPC variants such as KPC-35, highlighting the potential for emerging resistance even within this group (101).

Aztreonam/avibactam exhibited the broadest activity spectrum, showing full activity against all carbapenemase classes and combinations thereof. These findings are consistent with prior studies demonstrating 100% susceptibility of AZA against MBL, serine carbapenemase, and co-producing isolates (35, 36, 37). Such results underscore the therapeutic potential of AZA as a key agent for managing MDR infections, particularly those mediated by MBL-producing *Enterobacterales* (38). This is especially relevant in Peru, where NDM-type carbapenemases remain the most prevalent (4).

Cefiderocol represents a valuable second-line option against NDM and other MBL-producing *Enterobacterales*, with evidence supporting high clinical efficacy (39). In our study, all isolates—including co-producers—remained susceptible to cefiderocol (100%). Nevertheless, further research is warranted, as a recent meta-analysis identified multiple β-lactamases and resistance determinants associated with increased cefiderocol MICs, particularly among NDM-producing strains (40).

This study confirms that NDM-type MBLs remain the most prevalent carbapenemases in Peru, surpassing KPC variants (4). These findings position Peru among the countries where NDM represents the dominant carbapenemase, in line with global trends reporting its dissemination across all continents (41, 42).

In contrast, this pattern is not universal. Surveillance reports from Europe, Latin America, and the Caribbean indicate that KPC-type carbapenemases remain the most widespread among *Enterobacterales* in those regions, having reached endemic levels in several countries (41, 43, 44).

Historically, *K. pneumoniae* has been the main reservoir for carbapenemases, as reflected in this study (21). However, *E. coli* has shown a progressive increase in the dissemination of carbapenemase genes, particularly since 2015 (43). This observation mirrors the global trend of expanding MBL-type enzymes into species already associated with carbapenemase propagation (21, 43).

A distinctive feature of our region—also evident in this study—is the low prevalence of OXA-48-like carbapenemases (41). In our cohort, OXA-48-like enzymes accounted for 4.6% of isolates, showing only a modest increase compared with previous surveillance periods (4), ranking as the third most frequent carbapenemase detected among *Enterobacterales* in Peru. Interestingly, within *E. coli*, OXA-48-like was the second most common enzyme (31.1% in *E. coli* vs. 5.4% in *K. pneumoniae*).

An additional emerging concern is the increasing detection of co-producing carbapenemase isolates since the onset of the COVID-19 pandemic. *E. coli*, previously regarded as an occasional reservoir, has begun to harbor multiple carbapenemase genes more frequently (21). Our findings confirm the persistent circulation of *K. pneumoniae* co-producing KPC and NDM (3.1%) after the first sporadic case reported in 2016 (45), with a notable rise during the pandemic period (46). Moreover, the first *E. coli* isolate co-harboring *bla_NDM_* and *bla_OXA-48-like_* was reported in 2021, followed by subsequent detections with similar genetic configurations (47).

A recent review suggests that the emergence of multi-carbapenemase producers may be linked to the accumulation of non-β-lactam resistance mechanisms, likely driven by selective pressure from other antimicrobial classes (48).

Whole-genome sequencing (WGS) enabled the characterization of the allelic variants of carbapenemase genes previously identified by PCR. The most frequent variant detected was *bla*_NDM-1_ consistent with multiple reports from Peru (49, 50, 51). The *bla_KPC-2_* variant, also previously documented in the country (52), and *bla_OXA-48-_,* first reported in Peru in 2022 (53), were likewise identified. These variants represent the most commonly reported carbapenemase alleles across Latin America (41, 43).

Notably, *bla*_OXA-181_ was also detected—this variant has been recently reported in Peru among *K. pneumoniae*, *E. coli*, and *Citrobacter* spp. isolates (54), suggesting that its presence in the country is not sporadic but rather indicative of ongoing local dissemination. Additionally, *bla*_NDM-5_ was identified; this allele has been described in Argentina, Uruguay, Brazil, Mexico, United States, Europe and Middle east (55, 56, 57, 58, 59, 60, 61, 62). To our knowledge, this represents the first report of *bla*_NDM-5_ in Peru. Furthermore, co-producing isolates harboring *bla*_KPC-2_ + *bla*_NDM-1_ and *bla*_NDM-1_ + *bla*_OXA-48_ were detected, a phenomenon considered a global public health emergency (48).

Bioinformatic analysis of WGS data provided additional genomic insights into *E. coli* and *K. pneumoniae* isolates. MLST revealed two predominant *E. coli* clones: ST410 and ST167, which together accounted for 78.3% of sequenced *E. coli* isolates. *E. coli* ST410, belonging to clonal complex CC10, has been increasingly associated with the global spread of *bla*_OXA-181_ (48) and is now recognized as one of the most prevalent carbapenem-resistant *E. coli* (CRE) lineages worldwide, with presence across all five continents (63). In this study, the fimH24 variant of *E. coli* ST410 was the second most frequent sequence type (30%, 7/23), strongly associated with chromosomally located *bla*_OXA-48_. This finding is clinically relevant, as the presence of a single chromosomal copy of *bla*_OXA-48_ may hinder detection by standard phenotypic methods, posing an additional diagnostic challenge for clinical laboratories.

Regarding *E. coli* ST167, this lineage is among the most globally distributed and has shown increasing occurrence in Latin America. It is frequently associated with *bla*_NDM-5_, typically carried on IncF plasmids (63). In our study, ST167 was the predominant lineage among *E. coli*-CRE isolates (48%, 11/23), linked to the O101:H10 serotype and *bla*_NDM-5_ carriage. However, unlike previous reports, our isolates lacked the fimH54 allele, previously proposed as a conserved feature in this lineage conferring enhanced colonization and evolutionary advantages (64). This pattern supports the local spread of a closely related clonal group (6–21 SNPs apart) across several healthcare institutions in Peru.

Among *K. pneumoniae*, four major STs were identified—ST147, ST273, ST15, and ST45—representing 58.3% of sequenced isolates. Multidrug-resistant *K. pneumoniae* is recognized as a critical global health threat (1) and its rapid dissemination is largely attributed to the geographic expansion of successful clonal groups (CGs) such as CG15, CG101, CG147, CG258, and CG307 (65, 66). *K. pneumoniae ST147* has been identified as a globally distributed high-risk clone (67), closely related to ST273 and ST392, all of which belong to CG147 (68, 69). These clones commonly harbor fluoroquinolone-resistance mutations (*gyrA* S83I, *parC* S80I).

In our isolates, ST147 carried capsular loci KL64 (except one with KL20), while ST273 carried KL74. Historically, CG147 (primarily ST147) has been associated with multiple carbapenemase types since its emergence in the early 1990s, with ST273 first reported in 1995 (67, 70). Between 2010 and 2014, CG147 clones were reported globally in association with KPC, NDM, and OXA-48-like enzymes (71). In Peru, ST147 has been repeatedly identified and is emerging as a dominant clone responsible for carbapenemase dissemination in the country (72, 73, 74, 75).

*K. pneumoniae* ST15 is an emerging high-risk clone frequently implicated in hospital outbreaks (76). ST15 has been reported with multiple carbapenemases—including OXA-232, KPC-2, and NDM—in various settings (77) and has also been documented in Peru carrying NDM (78). In our series, ST15 isolates harbored QRDR mutations (*gyrA* S83I/D87A; *parC* S80I), the predominant capsular locus KL48, and the ybt1/ICEKp4 virulence module linked to siderophore-mediated iron acquisition (79, 80, 81).

*K. pneumoniae* ST45 has been described worldwide as a carrier of ESBLs and carbapenemases (82, 83, 84) and was recently reported in Peru as an NDM-producing lineage (78). In our 2020–2022 collection, ST45 isolates co-produced KPC-2 and NDM-1, a concerning pattern that intensified during the COVID-19 pandemic. These ST45 isolates lacked QRDR mutations, carried KL62, and encoded ybt1/ICEKp4, mirroring the virulence profile of our ST15 isolates.

Virulence in *K. pneumoniae* is associated with additional siderophores (*ybt, iuc, iro*) and specific capsular serotypes (K1, K2, K5) (85). Most hvKP infections arise in the Asia-Pacific region and are enriched in STs such as ST23, ST65, and ST86 (86). We identified one ST23 isolate carrying hallmark hvKP determinants (e.g., *iro, clb*) and rmpA associated with the hypermucoviscosity regulon. Importantly, hypermucoviscosity and hypervirulence are not synonymous—neither phenotype implies the other (87). Capsule overproduction (and thus mucoviscosity) can be modulated *by magA, rmpA, rmpA2*, and the two-component system *rcsAB* (87). Consistent with hvKP, our isolate belonged to capsular type K1 (86).

### Genetic context of carbapenemase genes

*bla*_NDM-1_.-All *bla*_NDM-1_ positive isolates shared a common backbone (ISAba125 - *bla*_NDM-1_ - *ble*_MBL_ -*trpF – dsbC – dct - groES*). This structure matches Peruvian plasmids IncFIB–IncHI1B pKpCol17ndm from a 2017 urine isolate (GenBank CP072906) (88) and pNDM1_Isoform5 from a 2016 isolate (GenBank MN816233.19) (89), and closely resembles Latin-American IncC type 1 contexts (pKQN17277, Argentina 2014, GenBank MH995507.1) (90) and pCf638 (Uruguay 2013, GenBank MT897966.1) (91). In our dataset, *bla*_NDM-1_ was carried either by hybrid IncFIB–IncHI1B plasmids (predominantly in *K. pneumoniae*) or IncC type 2 plasmids (present in both species). These plasmids co-harbored PMQR genes (*qnrE2, qnrVC1*), aminoglycoside acetyltransferases (*aac(3′)IIe*, *aac(6′)Ian*), and ESBLs (*bla*_TEM-26_, *bla*_PER-2_)—a resistance constellation frequently reported in South America (92, 93, 94).

*bla*_NDM-5_.-All *bla*_NDM-5_ isolates shared IS*Aba125* - *bla*_NDM-5_ – *ble*MBL – *trpF - dsbD*, identical to plasmids p32A19001_A_NDM (Canada 2019; GenBank PV023112.1) (95) and pEC26-NDM-5 (Bangladesh 2021; GenBank LC807790.1). *bla*_NDM-5_ occurred on IncFIA/IncFIB/IncFII plasmids and, almost uniformly, in ST167, a globally disseminated lineage (96, 97). Reports warning of emerging cefiderocol non-susceptibility among *bla*_NDM-5_ carriers warrant vigilance (98, 99).

*bla*_KPC-2_.- This gene resided within a non-Tn4401 element (*bla*_NKPC-1C_) featuring a partial IS*Kpn6* with its left inverted repeat and a Tn3 resolvase (*tnpR*) flanking *bla*_NKPC-2_. This arrangement is identical to pKP13d (Brazil 2014; GenBank CP003997.1) (100) and highly similar to pKP38_5 from Peru (GenBank CP159931.1) (101). The absence of Tn4401 repeats suggests recombination-mediated mobility via IS*Kpn6* and Tn3-family sequences (102, 103, 104). Most *bla*_KPC-2_ loci (10/15) were carried on IncX3–IncU plasmids, previously reported in clinical, colonization, and even food isolates in the region (105, 106).

*bla*_OXA-48_.- In our cohort, *bla*_OXA-48_ was chromosomally located and flanked by IS*1R* elements in a configuration resembling Tn2696 embedded within Tn1999 variant 3 (GenBank HE617182.1) (107). Nearly identical chromosomal contexts have been described in *E. coli* (New Zealand 2020, CP187211.1; Japan 2019, AP024694.1) and *K. pneumoniae* (China 2019, CP134036.1).

This analysis covers a restricted sampling window (four institutions in Lima, 2020–2023) and therefore may not capture the national picture. Long-read plasmid reconstruction was performed on a subset of representative isolates, providing partial but sufficient evidence that a limited set of successful plasmids is driving the dissemination of carbapenemases across lineages, hospitals, and years.

## CONCLUSIONS

Our WGS analysis of 83 CPE isolates from Peru documents the emergence and spread of high-risk *K. pneumoniae* lineages (ST147, ST15, ST45) and high-risk *E. coli* lineages (ST167, ST410), and—to our knowledge—the first detection of *bla*_NDM-5_ in the country. The recurrent association of epidemiologically fit plasmids with additional AMR determinants underscores the urgent need for coordinated antimicrobial stewardship, genomic surveillance, and infection-prevention programs to curb carbapenem resistance in Peru and the wider Latin-American region.

## ACKNOWLEDGMENTS

We thank the Faculty of Human Medicine, University of Piura, for funding this project (grant PI2307). We also acknowledge Pfizer Inc. for providing avibactam used in this study. Pfizer had no role in the study design, data collection, data interpretation, or decision to publish the results. The authors are grateful to Jose Matta-Chuquisapon, Christian Rivas, and Brenda Moy for their valuable technical assistance during this work.

## AUTHOR CONTRIBUTIONS

A.G.R. and E.G.E. conceived and designed the study. E.G.E., R.G.C.M., and L.A. performed laboratory experiments. J.C.G.T., E.S.C., G.P.L., C.J., M.M.C and R.S.A. contributed to data collection. A.G.R and E.G.E. performed genomic analysis. E.G.E and A.G.R. supervised the study and drafted the manuscript. All authors reviewed and approved the final version.

